# Transcranial static magnetic field stimulation over the left dorsolateral prefrontal cortex modulates EEG spectral activity and phase synchronization without detectable changes in divergent thinking

**DOI:** 10.64898/2026.11.18.767402

**Authors:** Eva Díez-Rodríguez, Daniel Jimeno-Huete, Antonio Oliviero, Vanesa Soto-León

## Abstract

**Background:** Transcranial static magnetic field stimulation (tSMS) can modulate cortical excitability, but its prefrontal electrophysiological effects remain incompletely characterized.

**Objective:** To investigate whether tSMS over the left dorsolateral prefrontal cortex (DLPFC) affects divergent-thinking performance and modulates EEG spectral activity and phase synchronization.

**Methods:** Forty-one healthy adults were randomly assigned to real (n = 21) or sham (n = 20) tSMS in a double-blind, sham-controlled study. Divergent thinking was assessed using the Alternative Uses Task (AUT) before and after stimulation. EEG was recorded at baseline, throughout 30 minutes of stimulation at rest, and during the subsequent AUT. Conventional spectral power, aperiodic parameters, and intra- and interregional phase-locking value (PLV) were analyzed using baseline-adjusted statistical models, with Holm correction for EEG contrasts.

**Results:** Divergent-thinking performance did not differ significantly between groups. During stimulation, real tSMS was associated with greater left frontal theta power in the final stimulation block, increased right frontal intra-regional theta synchronization, and greater beta and low-gamma bands phase synchronization involving frontal and centroparietal recording regions. During the subsequent AUT, greater bilateral frontal theta power, right frontal alpha power, right frontal aperiodic offset, and right frontal theta synchronization were observed following real stimulation.

**Conclusions:** A single session of left DLPFC tSMS can modulate frontal spectral activity and phase synchronization across distributed recording regions without significantly improving divergent-thinking performance. These findings highlight the electrophysiological responsiveness to prefrontal tSMS, although the functional significance of these changes remains to be established.

## 1. Introduction

Transcranial static magnetic field stimulation (tSMS) is an emerging non-invasive brain stimulation (NIBS) technique in which a neodymium magnet placed on the scalp generates a static magnetic field capable of modulating cortical excitability [1–3]. Since the initial demonstration that tSMS can reduce motor cortical excitability [1], subsequent studies have reported changes in neural excitability, oscillatory activity, and behavior across motor, somatosensory, visual, and supplementary motor regions [4–9]. However, the effects of tSMS over prefrontal regions remain insufficiently characterized. The left dorsolateral prefrontal cortex (DLPFC) is a major NIBS target [10] involved in working memory, cognitive flexibility, inhibitory control, and goal-directed behavior [11–13]. Only a few studies have examined tSMS over the left DLPFC, with mixed preliminary findings across behavioral and electrophysiological outcomes [14–18]. Whether left DLPFC tSMS modulates higher-order cognitive performance and how it influences prefrontal electrophysiological activity therefore remain open questions.

Divergent thinking provides a theoretically relevant behavioral domain for investigating the cognitive effects of left DLPFC stimulation. It refers to the generation of multiple possible responses to open-ended problems and is widely considered an indicator of creative potential [19,20]. This ability relies on executive and semantic processes, including cognitive flexibility, controlled retrieval, response selection, and the inhibition of dominant responses [21,22], several of which engage the DLPFC. Neuroimaging studies implicate the left DLPFC and distributed executive-control regions in creative idea generation [23–26], while NIBS studies suggest that stimulation of this region can influence divergent-thinking performance [27–30]. Creative ideation has also been associated with distinct electrophysiological patterns, including changes in alpha-band power during divergent-thinking tasks [31–33], functional coupling [34,35], and aperiodic activity [36]. The Alternative Uses Task (AUT), in which participants generate unusual uses for common objects, is a well-established measure of verbal divergent thinking that provides indices of fluency, flexibility, and originality [19,20]. Given the involvement of the left DLPFC in executive processes relevant to creative ideation, we used the AUT to assess the effects of prefrontal tSMS on verbal divergent-thinking performance and to explore associations between task performance and EEG activity.

Previous EEG studies have demonstrated that tSMS can modulate spectral power, although the affected frequencies and scalp regions may depend on the stimulation target. Studies applying tSMS over the occipital and parietal cortices have primarily reported increases in alpha-band power in regions associated with the stimulation site [6,7]. However, stimulation over the primary motor cortex (M1) has also been shown to increase theta-band power and interregional phase synchronization in the theta band [37]. The limited electrophysiological evidence from DLPFC-tSMS studies suggests heterogeneous findings, including exploratory regional patterns in alpha-, theta-, and beta-band power [14] and changes in task-related beta-band synchronization [18]. These findings motivate examining EEG spectral power across different frequency bands and scalp regions.

Because conventional band-limited power includes both periodic and aperiodic contributions, spectral parameterization can provide complementary information about the underlying spectral characteristics [38,39]. This approach allows characterization of the aperiodic background, which follows an approximately 1/f-like relationship. Its exponent describes the slope of this component and has been proposed as an indirect indicator of population-level excitation–inhibition dynamics, whereas the offset reflects its overall elevation [40,41]. The relevance of examining these parameters is further supported by evidence that other NIBS techniques can modulate aperiodic EEG activity [42–44]. In addition, the spatially distributed EEG patterns previously reported following DLPFC tSMS raise the question of whether phase relationships within and between scalp regions are also affected. Phase-locking value (PLV) quantifies the consistency of phase differences between EEG signals, providing a measure of intra- and interregional phase synchronization at the sensor level [45,46]. Together, these complementary EEG measures may help characterize the electrophysiological effects of prefrontal tSMS beyond changes in conventional spectral power alone.

We therefore aimed to characterize the electrophysiological effects of tSMS over the left DLPFC and to investigate whether stimulation modulates verbal divergent-thinking performance. To examine the EEG response over time, we recorded EEG throughout 30 minutes of stimulation, extending the 10-minute recording period used by Sheffield et al. [14] to investigate whether similar spectral patterns emerged and how they evolved over the longer period. Beyond conventional spectral power, we examined aperiodic activity and intra- and interregional phase synchronization, measures that remain largely unexplored in DLPFC-tSMS research. EEG was also recorded during the subsequent AUT, allowing us to characterize electrophysiological activity during divergent thinking and explore whether EEG measures obtained during stimulation and the post-stimulation task were associated with behavioral performance. Based on previous electrophysiological findings, we hypothesized that EEG spectral power would differ between real and sham tSMS. Changes in aperiodic activity and phase synchronization, as well as their potential associations with behavioral performance, were investigated as additional exploratory outcomes. Given the involvement of the left DLPFC in executive processes that support divergent thinking [21–26], we hypothesized that real tSMS over this region would influence subsequent AUT performance relative to sham stimulation. However, the direction of this effect was difficult to predict given the complex role of the DLPFC in creative idea generation, which relies on the interplay between executive control and flexible associative processing [21,22].

## 2. Methods

### 2.1. Participants

A total of 41 healthy volunteers (35 self-reported female; mean age = 31.71 years, SD = 10.45, range = 20–53 years) participated in the study. Thirty-seven participants were right-handed and four were left-handed. Inclusion criteria required participants to be adults without neurological or psychiatric disorders, substance abuse history, pregnancy, or implanted electronic or metallic devices. Participants were recruited via advertisement and provided written informed consent. The study was approved by the Ethics Committee of the Hospital Nacional de Parapléjicos of Toledo (approval no. CEIC 977) and conducted in accordance with the Declaration of Helsinki.

### 2.2. Experimental design and procedure

The study employed a randomized, double-blind, sham-controlled, between-subjects design, with participants assigned to real (n = 21) or sham (n = 20) stimulation. EEG preprocessing, feature extraction, and statistical analyses were conducted blind to stimulation condition. Sessions lasted approximately 80 min. After questionnaires and baseline AUT, continuous EEG was recorded during resting baseline, 30 min of real or sham tSMS over the left DLPFC, and post-stimulation AUT (Fig. 1A). Only the eyes-open resting segment was used as the pre-stimulation EEG baseline (T1).

**Figure 1.**
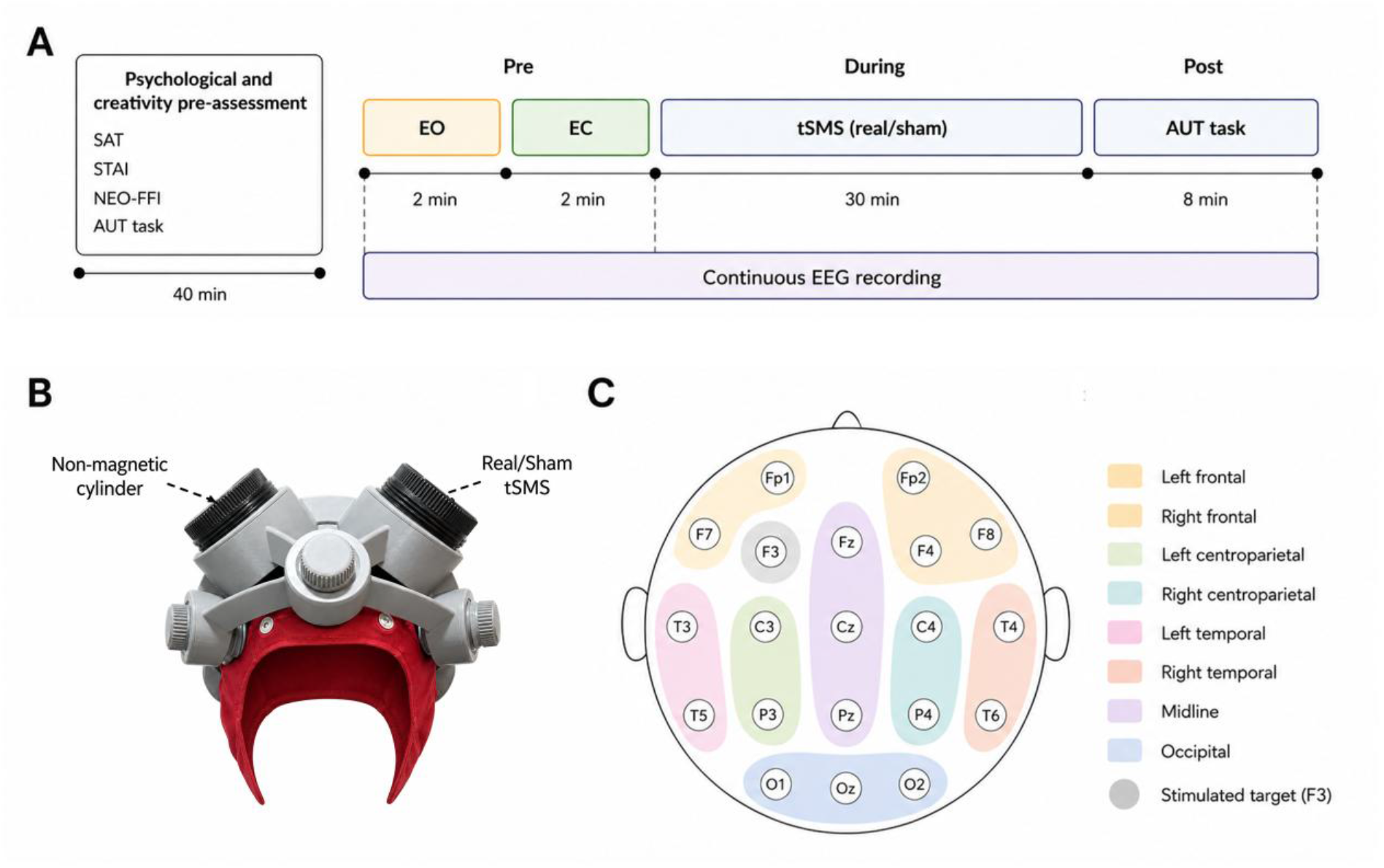
Experimental procedure and stimulation/EEG setup. **(A)** Experimental timeline. Participants completed psychological questionnaires and the pre-stimulation Alternative Uses Task (AUT) during the initial assessment period (40 min), followed by continuous EEG recording during baseline (2 min eyes open (EO), 2 min eyes closed (EC)), 30 min of real/sham tSMS, and the post-stimulation AUT (8 min). **(B)** tSMS setup showing the stimulation helmet used to secure the cylinders, with the real/sham tSMS cylinder positioned over F3 and the non-magnetic cylinder positioned over F4. **(C)** EEG electrode layout based on the international 10–20 system. Electrodes were grouped into regions of interest (ROIs), with frontal, centroparietal, and temporal regions subdivided by hemisphere, and midline and occipital electrodes treated as separate clusters. The grey circle indicates the stimulated target (F3).

### 2.3. Blinding integrity

Blinding integrity was assessed after the session by asking participants whether they had received real stimulation, sham stimulation, or did not know. Fifteen participants (36.6%) selected “did not know”. Of the remaining 26 participants, 14 correctly identified the stimulation condition (53.8%), with no significant association between perceived and received condition (Pearson’s χ²(1) = 0.16, p = .691).

### 2.4. Transcranial static magnetic field stimulation

Stimulation was applied over the left DLPFC using a cylindrical Ni–Cu–Ni-plated NdFeB magnet (60 mm diameter, 30 mm thickness, 370 g; model MAG60r; Neurek SL, Toledo, Spain), with a nominal force of 1,177 N. Previous modeling estimates a cortical magnetic field of approximately 120–200 mT at ∼1.2 cm depth [2,47]. Sham stimulation was delivered using a non-magnetic stainless-steel cylinder matched in size, weight, and appearance (model MAG60s; Neurek SL, Toledo, Spain). The real or sham cylinder was positioned over F3 according to the international 10–20 system and secured using an ergonomic helmet (model MAGdpv1.1; Neurek SL, Toledo, Spain; Fig. 1B). A nonmagnetic cylinder was placed over F4 as a counterweight.

### 2.5. Behavioral assessment

Divergent thinking was assessed using a computerized Alternative Uses Task (AUT). Participants were instructed to generate as many unusual uses as possible for common objects, with 2 min allocated to each object. Different sets of objects were administered before and after stimulation to minimize repetition and learning effects.

Responses were evaluated for fluency, flexibility, and originality. A composite AUT score was calculated by standardizing and averaging these measures. Originality was independently assessed by four raters blinded to stimulation condition, following established subjective and consensual-assessment approaches to divergent-thinking scoring [48,49], with good inter-rater reliability for averaged ratings (ICC = 0.87). Detailed scoring procedures are provided in Supplementary Methods.

Sustained attention, trait anxiety, and personality were assessed to characterize the sample and evaluate baseline group equivalence. These measures included a computerized sustained-attention task, the Spanish State–Trait Anxiety Inventory (STAI) [50], and the Spanish NEO Five-Factor Inventory (NEO-FFI) [51]. Additional details on the baseline measures are provided in Supplementary Methods.

### 2.6. EEG acquisition and preprocessing

EEG was recorded from 19 channels using a 21-channel Ag/AgCl cap positioned according to the international 10–20 system. The F3 electrode was excluded to optimize magnet-to-scalp contact over the stimulation target [2,14,47], and left-frontal activity was therefore characterized using the remaining electrodes within the corresponding region of interest (ROI). Fpz served as ground, signals were referenced online to the right earlobe, and data were acquired at a sampling rate of 512 Hz (Micromed S.p.A.), with an online 0.05-Hz high-pass filter and a 50-Hz notch filter. Electrode impedances were maintained below 10 kΩ.

EEG preprocessing was performed in MATLAB using EEGLAB [52] and custom scripts. Auxiliary channels were excluded, and continuous EEG was band-pass filtered between 0.5 and 40 Hz. Bad channels were identified using automated flatline, channel-correlation, and line-noise criteria and removed before applying Artifact Subspace Reconstruction (ASR) [53]. ASR was performed using a burst criterion of 20, with contaminated-segment rejection enabled.

Extended Infomax independent component analysis (ICA) [54] was subsequently performed, and artifactual components were identified and removed using ICLabel [55] with a probability threshold of 0.70. Previously rejected channels were then reconstructed by spherical interpolation.

EEG data were organized into five analysis windows: the pre-stimulation eyes-open baseline (T1), three consecutive 10-min stimulation blocks (T2: 0–10 min; T3: 10–20 min; T4: 20–30 min), and the post-stimulation AUT (T5). The original temporal structure was preserved during preprocessing, and spectral and connectivity analyses were performed on continuous EEG segments within each window. Additional preprocessing parameters are provided in the Supplementary Methods.

### 2.7. Spectral analysis

Power spectral density (PSD) was estimated using Welch’s method with 2-s Hamming windows, 50% overlap, and 0.5-Hz frequency resolution. Spectra were calculated separately for continuous EEG segments and combined using a weighted average based on the number of valid windows.

For conventional spectral analysis, outlying PSD values were identified using Tukey’s 1.5 × interquartile-range criterion in log10 power, followed by interpolation of isolated missing frequency bins. Absolute power was calculated for the theta (4–7.5 Hz), alpha (8–12.5 Hz), beta (13–29.5 Hz) and low-gamma (30–40 Hz) frequency bands and log10-transformed for statistical analysis.

The aperiodic component of the power spectrum was characterized using FOOOF (version 1.1.1) [38]. Original PSD estimates were parameterized over the 2–40 Hz frequency range using a fixed aperiodic model, yielding the aperiodic exponent and offset. Only fits meeting a predefined goodness-of-fit criterion (R² ≥ 0.70) were retained for subsequent analyses. Additional parameterization settings and quality-control procedures are described in the Supplementary Methods.

Spectral measures were averaged within eight predefined ROIs: left and right frontal, left and right centroparietal, left and right temporal, midline, and occipital regions (Fig. 1C). Regional power was averaged in linear units before log transformation, whereas aperiodic parameters were averaged across valid electrode-level estimates.

### 2.8. Phase synchronization

Phase synchronization was assessed using the phase-locking value (PLV) [45] in the theta, alpha, beta, and low-gamma frequency bands. Preprocessed EEG was band-pass filtered within each frequency band, and instantaneous phase was extracted from the analytic signal using the Hilbert transform.

PLV was calculated for each electrode pair from the phase differences across valid EEG samples. Intra-regional PLV was obtained by averaging synchronization values across all electrode pairs within each ROI. Inter-regional PLV was computed for all electrode pairs connecting two distinct ROIs, and the resulting pairwise PLV values were averaged to obtain a single connectivity estimate for each ROI pair.

Regional PLV estimates were Fisher z-transformed before statistical analysis. Additional filtering parameters, connectivity-estimation procedures, and quality-control criteria are described in the Supplementary Methods.

### 2.9. Statistical analysis

Statistical analyses were conducted in R version 4.5.2 [56]. Baseline group equivalence was assessed using independent-samples t-tests or Mann–Whitney U tests, as appropriate. All tests were two-tailed, with a significance threshold of α = .05.

Behavioral outcomes were analyzed using baseline-adjusted linear models, with post-stimulation performance as the dependent variable, Group (Real vs. Sham) as the factor of interest, and the corresponding pre-stimulation score as a covariate. AUT fluency, flexibility, originality, and the composite score were analyzed separately.

Electrophysiological outcomes were analyzed using Gaussian linear mixed-effects models fitted by restricted maximum likelihood (REML), with a participant-level random intercept and the corresponding pre-stimulation EEG value (T1) as a covariate. Models were fitted separately for each EEG measure and, where applicable, frequency band. Separate models were fitted for the stimulation period (T2–T4) and the post-stimulation AUT (T5). Stimulation-period models included Group, Time, ROI or Connection, and their interactions as fixed effects, whereas post-stimulation models included Group, ROI or Connection, and their interaction. Time was treated as a categorical factor.

Omnibus fixed effects were evaluated using Type III Wald chi-square tests. Estimated marginal means were used to obtain Real–Sham contrasts for each ROI or inter-regional connection, averaged across the stimulation period (T2–T4) and separately during the post-stimulation AUT (T5). Temporal effects were further examined through Group × Time interactions within each ROI or connection and block-specific Real–Sham contrasts. Additional within-group spatial and temporal comparisons were considered exploratory. P-values for planned and post hoc EEG contrasts were adjusted for multiple comparisons using the Holm procedure. All reported 95% confidence intervals were unadjusted for multiple comparisons.

Mixed-effects models were fitted using the R package glmmTMB, and estimated marginal means and contrasts were obtained using emmeans. Exploratory associations between electrophysiological measures and divergent-thinking performance were examined using regression models including Group, the corresponding EEG measure, and their interaction, adjusting for baseline AUT performance and the corresponding baseline EEG measure.

## 3. Results

### 3.1. Baseline characteristics and task equivalence

Baseline demographic, affective, personality, and AUT measures did not differ significantly between groups (Table 1).

**Table 1.** Baseline descriptive statistics by group.

| Variable | Sham | tSMS | *p* |
|---|---|---|---|
| Age (years) | 34.15 ± 11.99 | 29.38 ± 8.37 | .302 |
| Sustained attention RT | 279.43 ± 18.92 | 281.32 ± 31.56 | .949 |
| STAI–State | 16.50 ± 10.39 | 18.43 ± 8.15 | .511 |
| STAI–Trait | 18.65 ± 11.74 | 17.71 ± 8.56 | .979 |
| NEO-FFI Neuroticism | 18.65 ± 10.27 | 21.43 ± 9.88 | .383 |
| NEO-FFI Extraversion | 30.45 ± 9.51 | 32.67 ± 7.08 | .401 |
| NEO-FFI Openness | 28.30 ± 4.82 | 29.62 ± 6.11 | .449 |
| NEO-FFI Conscientiousness | 32.15 ± 5.92 | 31.81 ± 6.32 | .860 |
| NEO-FFI Agreeableness | 32.05 ± 8.75 | 33.67 ± 5.76 | .487 |
| AUT Fluency (Pre) | 6.37 ± 1.95 | 7.10 ± 2.52 | .358 |
| AUT Flexibility (Pre) | 4.81 ± 1.42 | 4.98 ± 1.21 | .687 |
| AUT Originality (Pre) | 2.04 ± 0.34 | 2.14 ± 0.36 | .377 |
| AUT Composite Index (Pre) | −0.13 ± 0.87 | 0.12 ± 0.85 | .375 |

*Note***.** Values are means ± SD. Sham group: n = 20; Real tSMS group: n = 21. AUT = Alternative Uses Task; STAI = State–Trait Anxiety Inventory; NEO-FFI = NEO Five-Factor Inventory.

### 3.2. Behavioral results: Alternative Uses Task

Baseline-adjusted analyses showed no significant between-group differences in post-stimulation AUT fluency, flexibility, originality, or composite score (all *p* ≥ .261; Table 2).

**Table 2.** Post-stimulation AUT descriptive statistics and baseline-adjusted group effects.

| Outcome | Sham | tSMS | t | df | *p* |
|---|---|---|---|---|---|
| Fluency | 6.12 ± 1.75 | 6.60 ± 1.82 | -0.265 | 36 | .793 |
| Flexibility | 4.58 ± 1.14 | 4.98 ± 1.12 | -1.141 | 36 | .261 |
| Originality | 1.93 ± 0.41 | 1.96 ± 0.40 | 0.465 | 36 | .644 |
| AUT composite | -0.13 ± 0.91 | 0.11 ± 0.85 | -0.393 | 36 | .696 |

*Note.* Group means and standard deviations describe the observed post-stimulation scores. The *t* statistics and two-sided *p* values refer to the group effect in linear models adjusted for the corresponding baseline score. AUT = Alternative Uses Task. tSMS = transcranial static magnetic field stimulation.

### 3.3. Spectral power

During stimulation (T2–T4), spectral power analyses revealed significant Group × ROI interactions in the theta (χ²(7) = 14.94, *p* = .037) and beta bands (χ²(7) = 22.13, *p* = .002), indicating that between-group differences varied across scalp regions. A significant Group × Time interaction was also observed in the theta band (χ²(2) = 6.78, *p* = .034). During the post-stimulation AUT (T5), significant Group × ROI interactions were observed in the theta (χ²(7) = 27.77, *p* < .001) and alpha bands (χ²(7) = 20.68, *p* = .004).

In the theta band, left frontal power averaged across the three 10-minute stimulation blocks (T2–T4) was estimated to be higher under real tSMS than sham (*z* = 2.54, 95% CI [0.032, 0.247], *p* = .011, *pHolm* = .090). Exploratory block-specific comparisons revealed significantly greater left frontal theta power under real stimulation during the final 10 minutes of stimulation (T4; *z* = 2.78, 95% CI [0.052, 0.303], *p* = .006, *pHolm* = .044; Fig. 2). During the same block, higher estimated theta power was also observed in the right frontal region (*z* = 2.24, 95% CI [0.018, 0.270], *p* = .025, *pHolm* = .157) and the midline region (*z* = 2.28, 95% CI [0.021, 0.274], *p* = .022, *pHolm* = .157), with neither comparison reaching significance after Holm correction.

**Figure 2.**
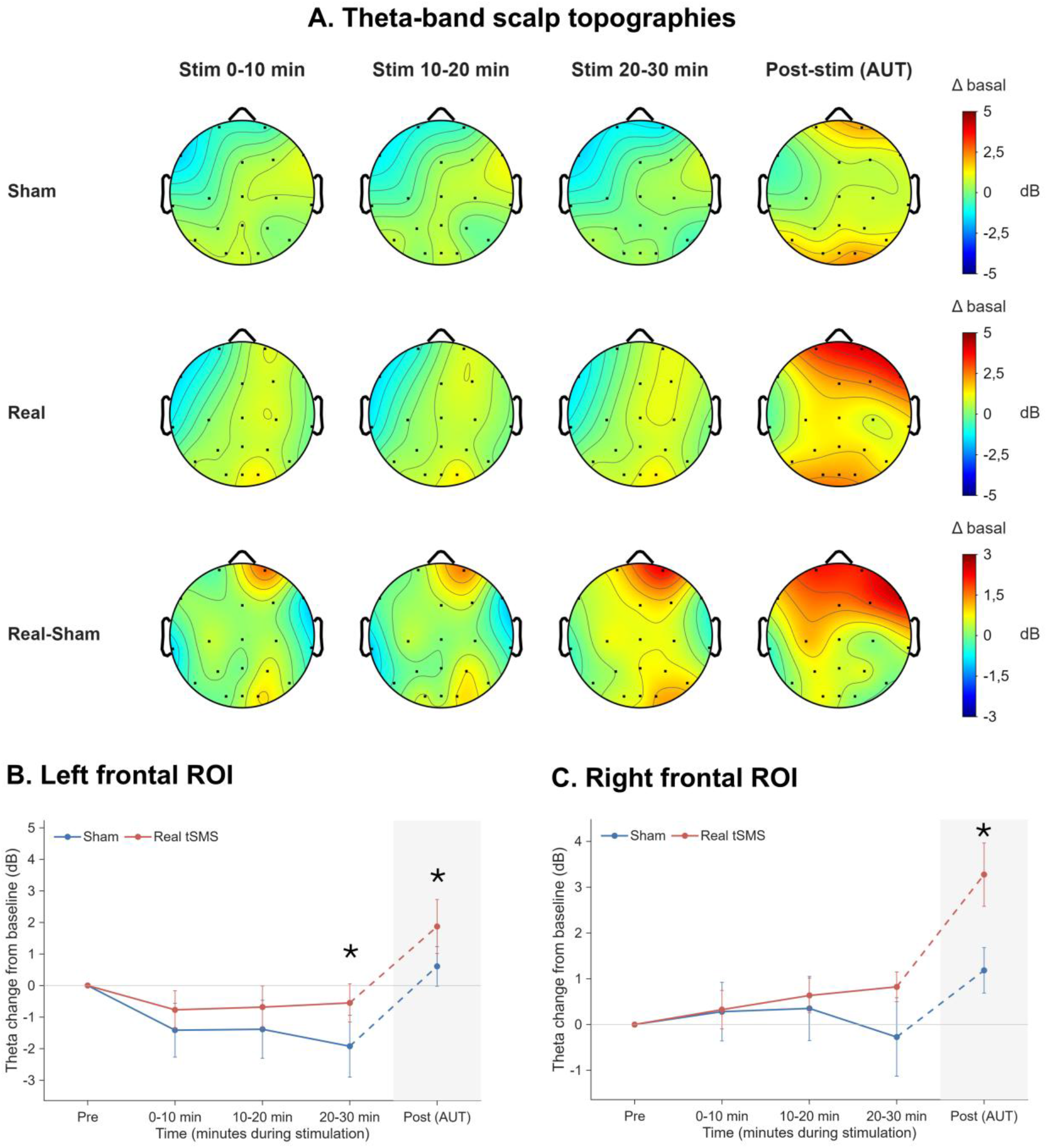
Theta-band power during stimulation and the subsequent Alternative Uses Task (AUT). **(A)** Scalp maps show mean changes from baseline across three 10-min stimulation blocks and the AUT. Color scales span ±5 dB for the sham and real tSMS groups and ±3 dB for their difference (real − sham). **(B–C)** Mean changes in the left (Fp1/F7) and right (Fp2/F4/F8) frontal regions, respectively. Error bars show ±1 standard error of the mean (SEM). Asterisks mark real–sham contrasts significant after Holm correction across eight regions (adjusted *p* < .05). Shading marks the AUT.

During the post-stimulation AUT (T5), theta-band power was significantly greater following real tSMS than sham stimulation in both the left frontal (*z* = 3.09, 95% CI [0.062, 0.277], *p* = .002, *pHolm* = .014) and right frontal regions (*z* = 4.57, 95% CI [0.145, 0.364], *p* < .001, *pHolm* < .001; Fig. 2). Higher estimated theta power was also observed in the midline region (*z* = 2.05, 95% CI [0.005, 0.226], *p* = .040, *pHolm* = .241).

In the beta band, left frontal power averaged across the 30-minute stimulation period (T2–T4) was estimated to be higher under real stimulation than sham (*z* = 2.06, 95% CI [0.005, 0.177], *p* = .039, *pHolm* = .313; Fig. 3). No regional beta-band contrast survived Holm correction.

**Figure 3.**
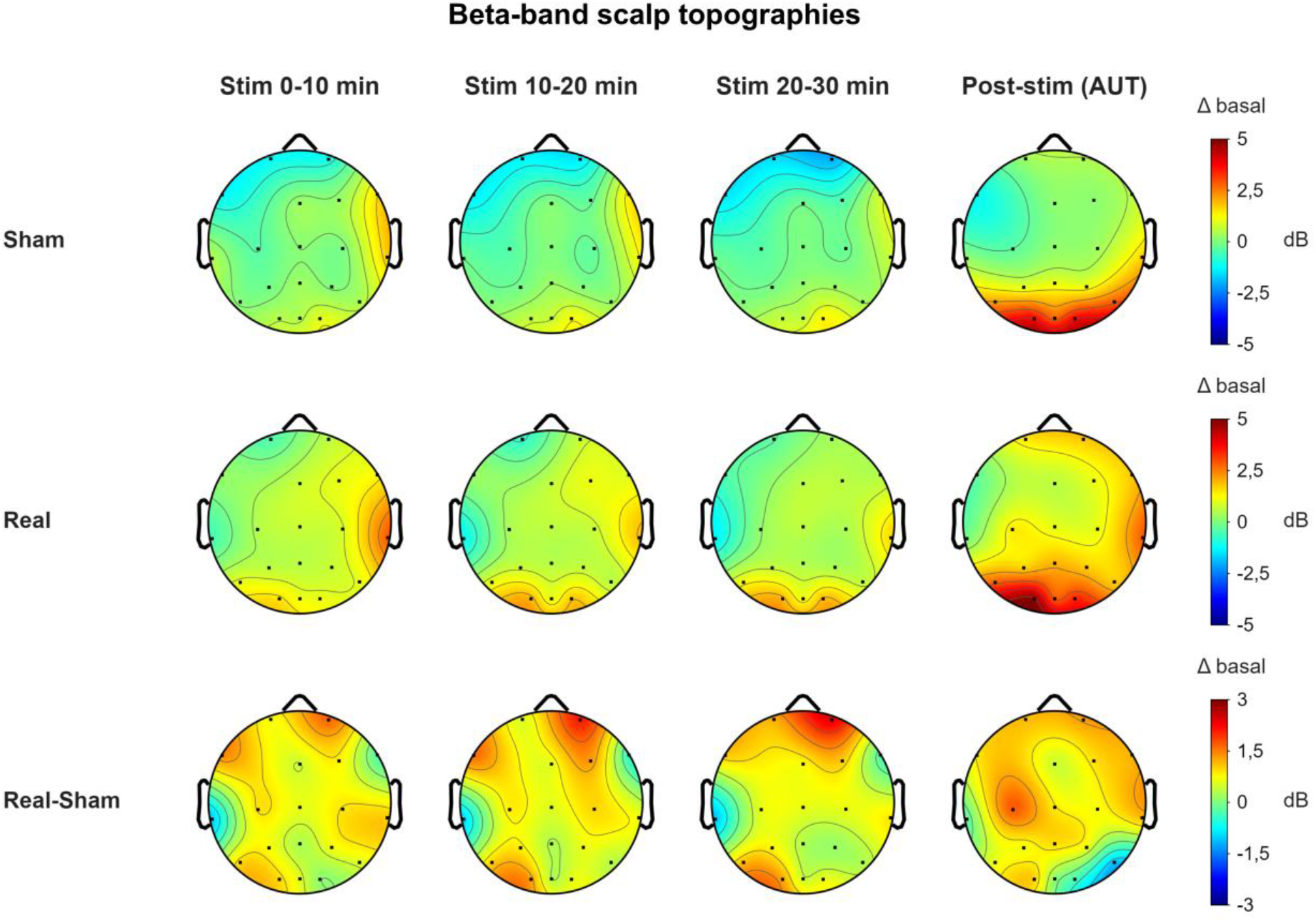
Beta-band power during stimulation and the subsequent Alternative Uses Task (AUT). Scalp maps show mean changes from baseline across three 10-min stimulation blocks and the AUT. From top to bottom, rows show the sham group, the real tSMS group, and their difference (real − sham). Color scales span ±5 dB for the group maps and ±3 dB for the difference maps. The maps are descriptive.

Finally, in the alpha band, power during the post-stimulation AUT (T5) was significantly greater in the right frontal region following real stimulation than sham (*z* = 3.55, 95% CI [0.087, 0.303], *p* < .001, *pHolm* = .003; Fig. 4).

**Figure 4.**
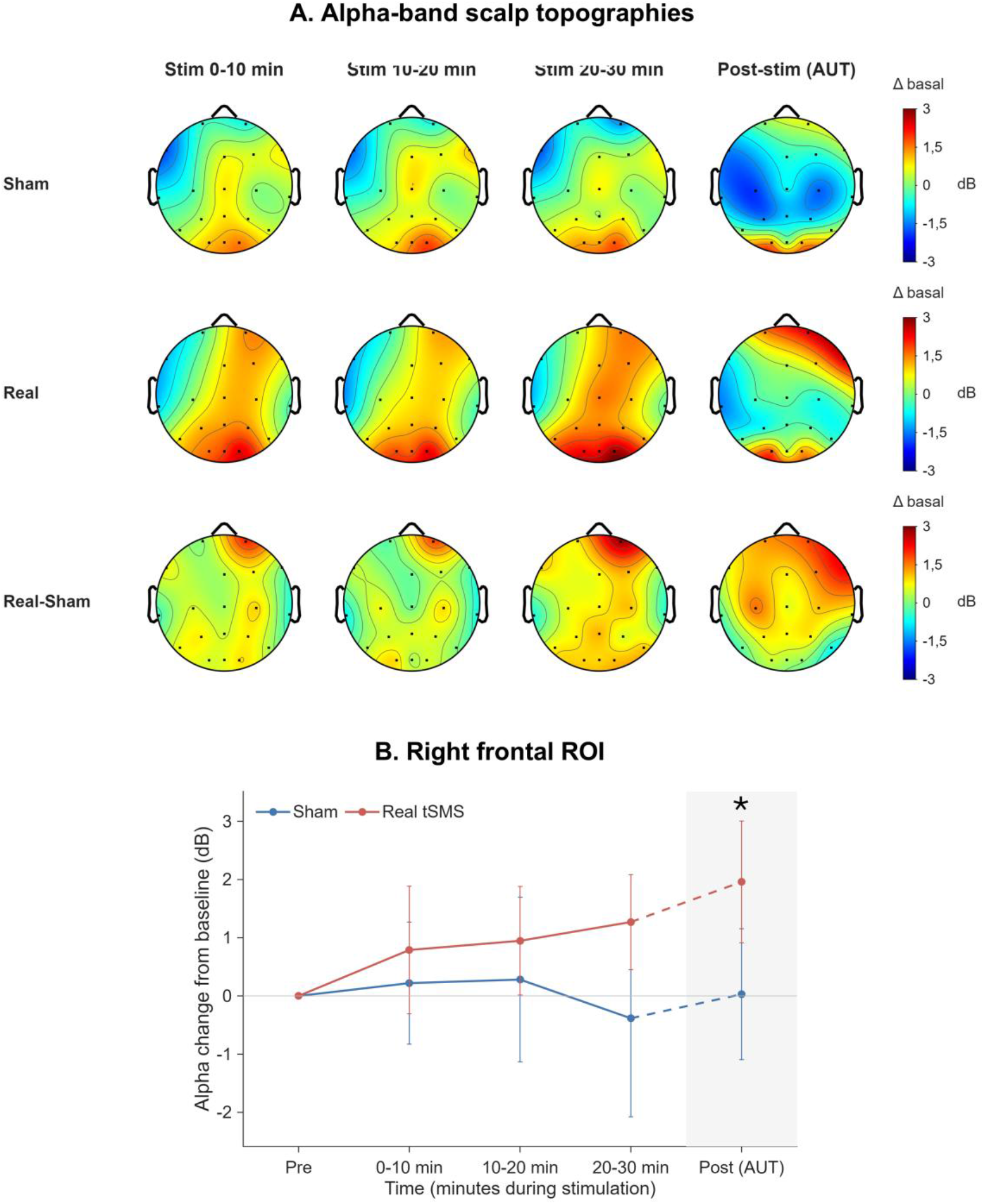
Alpha-band power during stimulation and the subsequent Alternative Uses Task (AUT). **(A)** Scalp maps show mean changes from baseline (dB) for the sham and real tSMS groups and their difference (real − sham) across three 10-min stimulation blocks and the AUT. **(B)** Mean change in the right frontal region (Fp2/F4/F8). Error bars show ±1 SEM. The asterisk marks the real–sham contrast during the AUT (shaded), which remained significant after Holm correction across eight regions (adjusted *p* < .05).

### 3.4. Aperiodic EEG activity

During stimulation (T2–T4), the aperiodic offset showed a significant Group × ROI interaction (χ²(7) = 18.02, *p* = .012), although no regional Real–Sham contrast averaged across the three stimulation blocks survived Holm correction. No significant Group × ROI interaction was observed for the aperiodic exponent during stimulation (χ²(7) = 7.76, *p* = .355).

During the post-stimulation AUT (T5), significant Group × ROI interactions were observed for both the aperiodic offset (χ²(7) = 26.54, *p* < .001) and exponent (χ²(7) = 14.70, *p* = .040). The offset was significantly higher following real tSMS than sham stimulation in the right frontal region (*z* = 3.08, 95% CI [0.149, 0.667], *p* = .002, *pHolm* = .016). Higher estimated offset was also observed in the left frontal region (*z* = 2.05, 95% CI [0.012, 0.530], *p* = .040, *pHolm* = .283).

The aperiodic exponent also showed higher estimated values in the right frontal region following real stimulation (*z* = 2.11, 95% CI [0.015, 0.425], *p* = .035, *pHolm* = .280). No regional Real–Sham contrast for the exponent survived Holm correction during either stimulation or the post-stimulation AUT.

### 3.5. Intra-regional phase synchronization

During stimulation (T2–T4), intra-regional theta-band phase-locking value (PLV) showed a significant Group × ROI interaction (χ²(7) = 14.34, *p* = .045). When averaged across the three stimulation blocks, regional comparisons revealed significantly greater theta-band synchronization within the right frontal region under real stimulation than sham (*z* = 2.86, 95% CI [0.057, 0.304], *p* = .004, *pHolm* = .034; Fig. 5).

**Figure 5.**
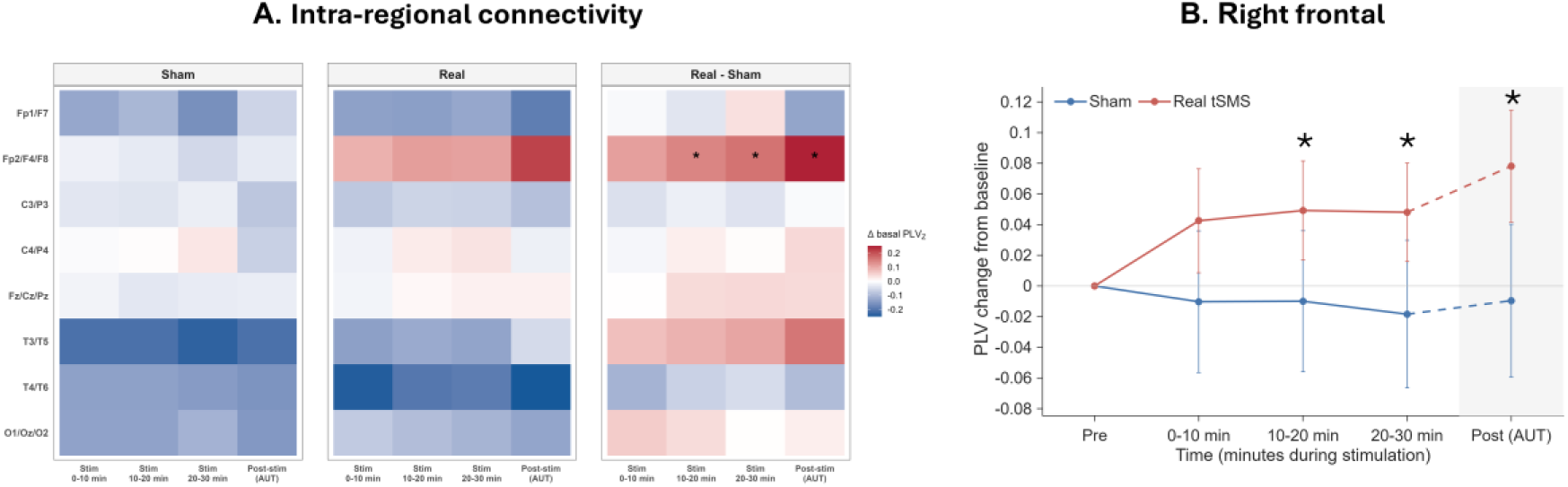
Intraregional theta-band phase synchronization. **(A)** Heatmaps show mean changes from baseline in Fisher z-transformed phase-locking value (PLV) within eight regions for the sham and real tSMS groups and their difference (real − sham). Columns represent three 10-min stimulation blocks and the subsequent Alternative Uses Task (AUT). **(B)** Mean change in right frontal PLV (Fp2/F4/F8). Error bars show ±1 SEM. Asterisks mark significant real–sham contrasts from baseline-adjusted mixed-effects models (Holm-adjusted *p* < .05). The contrasts for the stimulation-period average and the AUT were each corrected across eight regions. For regions with a significant stimulation-period contrast, follow-up contrasts were corrected across the three blocks. Shading marks the AUT.

Exploratory block-specific comparisons showed greater right frontal theta synchronization under real stimulation from 10 minutes onwards, with significant Real–Sham differences observed during T3 (*z* = 2.34, 95% CI [0.030, 0.343], *p* = .019, *pHolm* = .039) and T4 (*z* = 2.48, 95% CI [0.042, 0.358], *p* = .013, *pHolm* = .039; Fig. 5).

During the post-stimulation AUT (T5), although the overall Group × ROI interaction was not significant (χ²(7) = 13.38, *p* = .063), the ROI-specific Real–Sham comparison revealed significantly greater right frontal theta synchronization following real stimulation (*z* = 3.43, 95% CI [0.129, 0.475], *p* < .001, *pHolm* = .005; Fig. 5).

### 3.6. Inter-regional phase synchronization

During stimulation (T2–T4), inter-regional PLV analyses revealed significant Group × Connection interactions in both the beta (χ²(27) = 112.70, *p* < .001) and low-gamma bands (χ²(27) = 113.55, *p* < .001), indicating that between-group differences varied across inter-regional connections.

In the beta band, comparisons averaged across the three 10-minute stimulation blocks identified significantly greater synchronization under real stimulation between the left frontal and left centroparietal regions (*z* = 3.30, 95% CI [0.042, 0.164], *p* < .001, *pHolm* = .027) and between the right frontal and left centroparietal regions (*z* = 3.12, 95% CI [0.036, 0.159], *p* = .002, *pHolm* = .046; Fig. 6).

**Figure 6.**
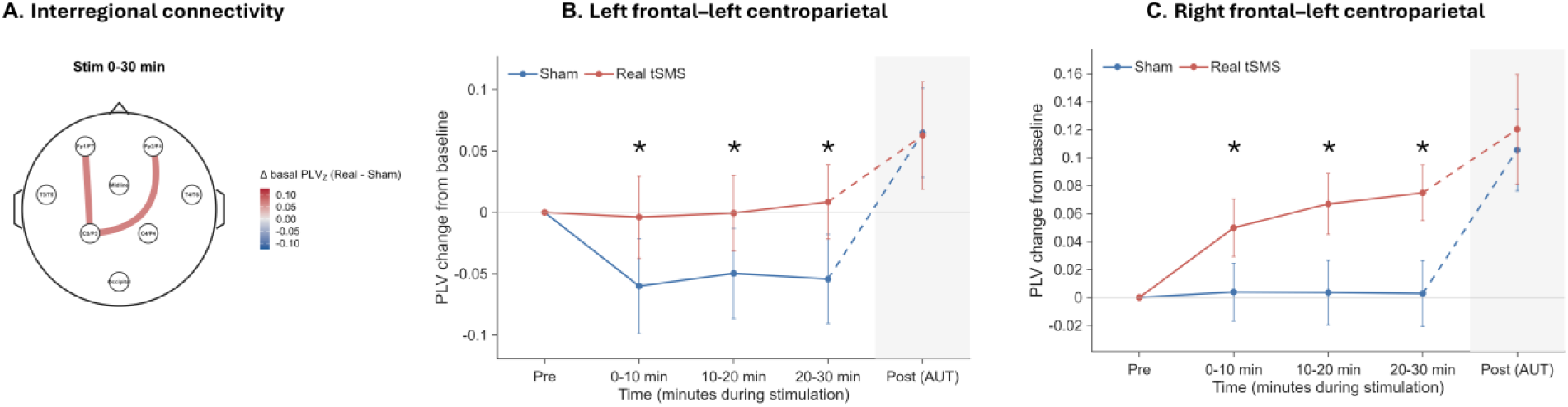
Interregional beta-band phase synchronization. **(A)** The map shows region pairs with a significant real–sham contrast averaged over 30 min of stimulation (Holm correction across 28 pairs). Line color and width represent the observed difference between groups in the change from baseline in Fisher z-transformed phase-locking value (PLV). Red indicates a positive real − sham difference in change from baseline. **(B–C)** Mean PLV changes between the left centroparietal region and the left **(B)** or right **(C)** frontal region. Error bars show ±1 SEM. Asterisks mark real–sham contrasts from baseline-adjusted mixed-effects models that remained significant after Holm correction across the three stimulation blocks within each pair (adjusted *p* < .05). Shading marks the subsequent Alternative Uses Task (AUT).

In the low-gamma band, mean PLV across T2–T4 was significantly greater under real stimulation in several frontal and frontoparietal connections. These included the left–right frontal connection (*z* = 3.25, 95% CI [0.045, 0.182], *p* = .0012, *pHolm* = .027), the left frontal–left centroparietal connection (*z* = 3.24, 95% CI [0.045, 0.182], *p* = .0012, *pHolm* = .027), and the right frontal–left centroparietal connection (*z* = 3.37, 95% CI [0.049, 0.187], *p* < .001, *pHolm* = .018; Fig. 7).

**Figure 7.**
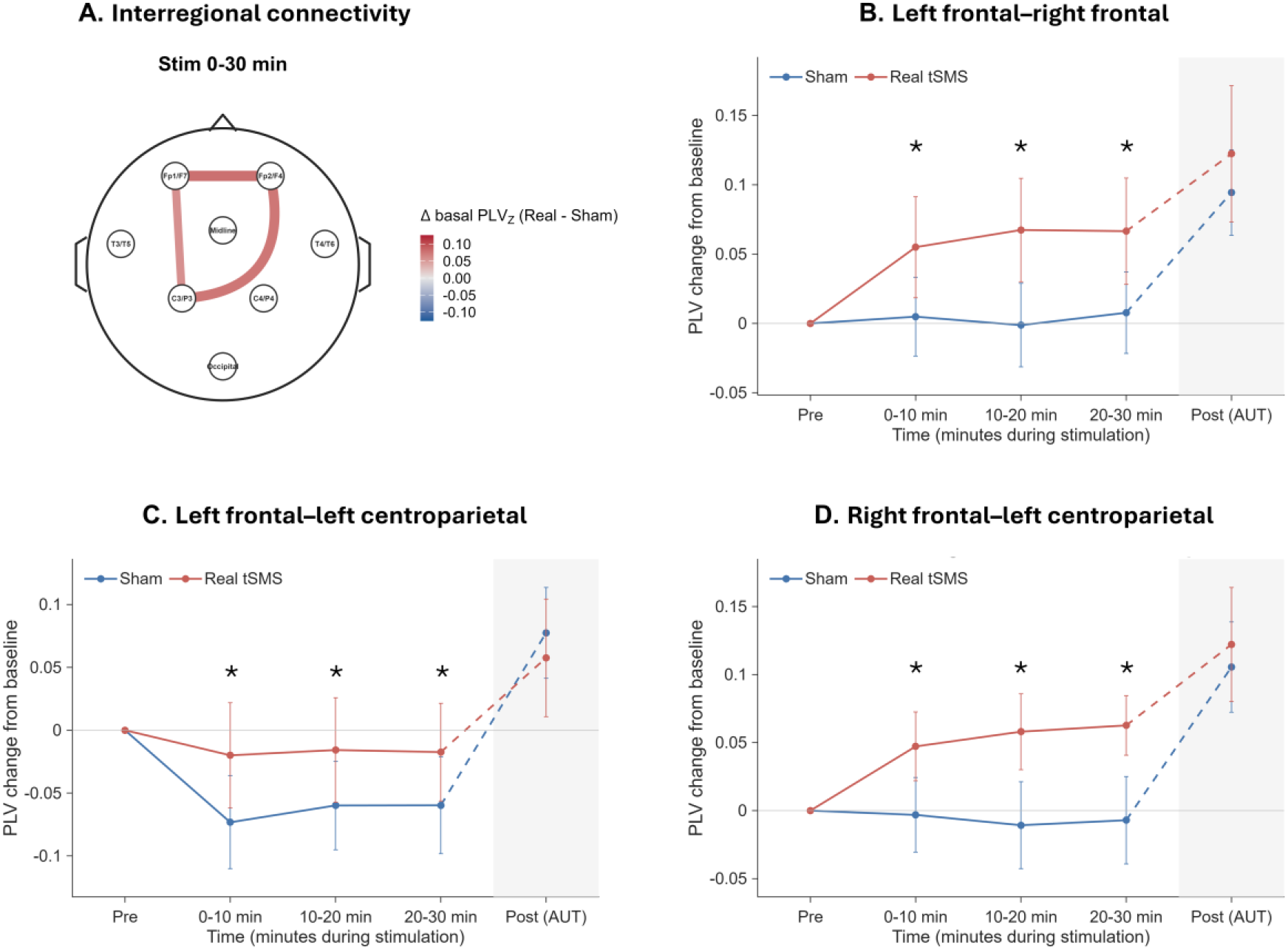
Interregional low-gamma-band phase synchronization. **(A)** The map shows region pairs with a significant real–sham contrast averaged over 30 min of stimulation (Holm correction across 28 pairs). Line color and width represent the observed difference between groups in the change from baseline in Fisher z-transformed phase-locking value (PLV). Red indicates a positive real − sham difference in change from baseline. **(B–D)** Mean PLV changes between the left and right frontal regions **(B)**, the left frontal and left centroparietal regions **(C)**, and the right frontal and left centroparietal regions (**D)**. Error bars show ±1 SEM. Asterisks mark real–sham contrasts from baseline-adjusted mixed-effects models that remained significant after Holm correction across the three stimulation blocks within each pair (adjusted *p* < .05). Shading marks the subsequent Alternative Uses Task (AUT).

Block-specific comparisons showed consistently positive estimated Real–Sham differences across T2, T3, and T4 in these beta and low-gamma band connections. The corresponding contrast statistics and Holm-corrected *p*-values for each stimulation block are presented in Table 3.

**Table 3.** Exploratory block-specific Real–Sham contrasts for selected inter-ROI PLV connections.

| Band | Connection | Block | z | *p<sub>Holm</sub>* |
|---|---|---|---|---|
| Beta | L. Frontal – L. Centroparietal | T2 | 2.594 | .0214 |
|  |  | T3 | 2.336 | .0214 |
|  |  | T4 | 2.691 | .0214 |
|  | R. Frontal – L. Centroparietal | T2 | 1.968 | .0491 |
|  |  | T3 | 2.485 | .0259 |
|  |  | T4 | 2.758 | .0174 |
| Low-gamma | L. Frontal – R. Frontal | T2 | 2.182 | .0291 |
|  |  | T3 | 2.731 | .0189 |
|  |  | T4 | 2.486 | .0258 |
|  | L. Frontal – L. Centroparietal | T2 | 2.668 | .0229 |
|  |  | T3 | 2.447 | .0288 |
|  |  | T4 | 2.260 | .0288 |
|  | R. Frontal – L. Centroparietal | T2 | 2.232 | .0256 |
|  |  | T3 | 2.710 | .0190 |
|  |  | T4 | 2.731 | .0190 |

*Note*. Values represent block-specific comparisons between Real and Sham tSMS during the three 10-minute stimulation blocks (T2, T3, and T4). Positive z-values indicate higher baseline-adjusted PLV in the Real tSMS group than in Sham. pHolm values were adjusted using the Holm procedure across the three stimulation blocks within each connection. L. Frontal = left frontal; R. Frontal = right frontal; L. Centroparietal = left centroparietal.

During the post-stimulation AUT (T5), no significant Group × Connection interactions were observed in either the beta (χ²(27) = 14.28, *p* = .978) or low-gamma band (χ²(27) = 12.65, *p* = .991). No individual inter-regional Real–Sham contrasts survived Holm correction during this phase.

### 3.7. Exploratory EEG–behavior associations

Exploratory EEG–behavior analyses suggested a group-dependent association between mean left frontal beta-band power during stimulation (T2–T4) and subsequent AUT fluency (*p* = .010), with opposite association slopes in the real and sham groups. Additional exploratory analyses suggested associations between right frontal intra-regional theta synchronization and divergent-thinking performance in the sham group, and between left frontoparietal beta synchronization and originality in the real stimulation group. None of these exploratory associations or interactions survived Holm correction.

## 4. Discussion

The present study shows that transcranial static magnetic field stimulation (tSMS) over the left dorsolateral prefrontal cortex (DLPFC) was associated with electrophysiological differences, without significant changes in divergent-thinking performance. Spectral power differences were predominantly frontal, involving theta activity during stimulation and theta and alpha activity during the subsequent Alternative Uses Task (AUT). A greater right frontal aperiodic offset was also observed during the post-stimulation AUT. In addition, real tSMS was associated with greater right frontal intra-regional theta synchronization during both phases and greater beta and low-gamma bands phase synchronization between frontal and centroparietal recording regions during stimulation.

Previous tSMS studies have reported heterogeneous electrophysiological responses across stimulation targets. Occipital and parietal tSMS has been associated primarily with increases in alpha-band power [6,7], whereas stimulation over the primary motor cortex has been shown to increase theta-band power [37]. Sheffield et al. [14], in turn, described exploratory patterns involving alpha, theta, and beta activity across different scalp regions following DLPFC stimulation. Interestingly, these same frequency bands were also implicated in our findings, although the significant spectral power differences were predominantly frontal, in contrast to the more spatially distributed patterns suggested by Sheffield et al. In our study, broader spatial involvement was instead apparent mainly through differences in phase synchronization between frontal and centroparietal recording regions. Previous studies have also reported changes in motor–parietal theta coupling following motor cortex tSMS [37] and reduced resting-state sensorimotor functional connectivity using fMRI [57]. Although these studies differ in their stimulation targets and measurement techniques, their findings suggest that tSMS may influence both regional activity and measures of interregional functional coupling.

Theta activity was particularly prominent in our findings. Greater left frontal theta power emerged during the final 10 minutes of stimulation, while greater theta power was observed in both frontal regions during the subsequent AUT. Theta oscillations have been associated with working memory, cognitive control, and the coordination of neural activity [58–60], raising the possibility that these differences involve neural processes relevant to higher-order cognition. However, the present findings do not allow these processes to be identified specifically. Real tSMS was also associated with greater right frontal alpha power during the AUT, a finding of interest given the relationship between alpha activity, internally directed attention, and creative ideation [31–35]. However, these associations are not specific to creativity, and greater theta or alpha power cannot be interpreted as enhanced cognitive processing. Left frontal beta power was also estimated to be higher during stimulation, although the difference did not survive Holm correction. Watanabe et al. [18] previously reported increased task-related beta synchronization following prefrontal tSMS, suggesting that beta-band activity may be sensitive to stimulation, although the measures and experimental conditions differed.

The spectral changes observed during the post-stimulation AUT may partly reflect the cognitive demands of the task. Descriptively, frontal theta and alpha power appeared to increase from the final stimulation block to the subsequent AUT in both groups. Previous EEG studies of divergent-thinking tasks, including the AUT, have reported increases in alpha power during idea generation, with alpha activity also associated with the originality of generated ideas [31–34]. Theta activity has also been linked to cognitive processes relevant to divergent thinking, including working memory and cognitive control [58–60]. Nevertheless, during the AUT, frontal theta and alpha power were greater following real than sham stimulation in specific regions, raising the possibility that prior stimulation influenced electrophysiological activity during subsequent task performance. However, this transition involved both removal of the magnet and initiation of active idea generation. Consequently, the present design cannot determine whether these differences reflect persisting stimulation-related effects, task-dependent responses, or their interaction. Furthermore, because EEG was not recorded during a matched post-stimulation resting condition, these findings cannot establish whether the observed differences were specifically related to divergent thinking. Future studies incorporating matched resting and task conditions during and after stimulation would help distinguish temporal from state-dependent responses.

Because conventional band-limited power contains both periodic and aperiodic contributions [38,39], we also examined whether the spectral differences were accompanied by changes in aperiodic activity. However, evidence for such changes was more limited. A significantly greater right frontal aperiodic offset was observed following real stimulation during the subsequent AUT, whereas the higher estimated exponent in this region did not survive Holm correction. Moreover, no regional aperiodic contrasts survived correction during stimulation. These findings therefore do not establish that the observed spectral power differences, particularly those emerging during stimulation, were driven by changes in the aperiodic component. Although a steeper aperiodic slope has been associated with greater relative inhibition or reduced cortical excitability [40,41], this relationship is not sufficiently specific to infer an inhibitory mechanism from our results. Similarly, differences in the aperiodic offset cannot be interpreted as direct evidence of changes in cortical excitability.

Phase synchronization provided complementary information. Real tSMS was associated with greater intra-regional theta synchronization in the right frontal region during stimulation and the subsequent AUT. Frontal theta synchronization has been linked to cognitive control and performance monitoring [61], although most evidence concerns task-related midfrontal activity. The present findings therefore raise the possibility that prefrontal tSMS influences frontal phase relationships, but their functional relevance remains to be established.

Inter-regional analyses revealed greater beta and low-gamma bands synchronization during real tSMS, particularly between frontal and centroparietal recording regions, with additional bilateral frontal involvement in low gamma. Greater synchronization under real tSMS was observed throughout the stimulation period, with significant between-group differences detected both when averaging across the three blocks and in block-specific analyses. However, the temporal trajectories appeared to differ across connections. In the left frontal–left centroparietal connection, beta and low-gamma synchronization tended to decrease from baseline under sham stimulation while remaining relatively stable under real tSMS. In contrast, synchronization tended to increase under real stimulation in the right frontal–left centroparietal connection and between bilateral frontal regions, while remaining closer to baseline under sham. Although descriptive, these patterns suggest that the influence of tSMS on inter-regional phase relationships may depend on the connections examined rather than reflecting a uniform increase in synchronization. No inter-regional beta or low-gamma bands differences survived correction during the subsequent AUT.

The involvement of frontal and centroparietal recording regions is particularly relevant given the established role of frontoparietal interactions in attention, working memory, and executive control, including the selection and maintenance of task-relevant information [62,63]. Interactions involving executive-control and default-mode networks have also been implicated in divergent thinking [21,22,25,26]. However, the greater frontocentroparietal phase synchronization observed during stimulation was not accompanied by improved subsequent AUT performance. Moreover, our sensor-level PLV estimates cannot be attributed to specific anatomical networks or cognitive processes, and greater synchronization does not necessarily indicate more efficient information processing. This is particularly relevant because the inter-regional differences occurred without concurrent cognitive demands. Studies combining prefrontal tSMS with executive and working-memory tasks could help establish whether these electrophysiological changes have functional consequences.

Despite the observed electrophysiological differences, real tSMS did not significantly affect AUT fluency, flexibility, originality, or composite performance compared with sham stimulation. This finding did not support our behavioral hypothesis and is consistent with the heterogeneous behavioral outcomes reported in previous studies of DLPFC tSMS [15–18]. One possible explanation is that a single session of tSMS may have been insufficient to produce detectable changes in divergent-thinking performance. Additionally, the AUT may have had limited sensitivity to subtle stimulation-related effects in this sample of healthy participants, whose behavioral performance showed limited variability. Furthermore, divergent thinking depends on interacting semantic, associative, and executive processes [21,22,25], and modulation of neural activity associated with one of these processes may not translate into measurable changes in overall task performance. Stimulation was also delivered at rest, whereas creativity was assessed afterward, and the relatively small sample may have limited statistical precision. These explanations remain speculative, and the absence of significant group differences should not be interpreted as evidence of behavioral equivalence.

Exploratory EEG–behavior analyses suggested that the relationship between left frontal beta power during stimulation and subsequent AUT fluency differed between real and sham conditions. Specifically, higher beta power tended to be associated with greater subsequent fluency under real stimulation, whereas the opposite pattern was observed under sham stimulation. However, these associations did not survive correction for multiple comparisons and should therefore be interpreted as preliminary, hypothesis-generating findings rather than evidence of a reliable relationship between electrophysiological activity and divergent-thinking performance.

These findings should be interpreted in light of methodological limitations. AUT performance provides a global and partly rater-dependent measure of divergent thinking that can be influenced by item characteristics, response strategies, and scoring procedures [20,48,49]. Future studies incorporating process-specific tasks could help determine whether tSMS affects particular components of divergent thinking. Because stimulation was delivered at rest and behavior was assessed afterward, further research should incorporate task-concurrent stimulation alongside resting-state protocols, with EEG recordings obtained under matched cognitive conditions. The EEG montage also imposed spatial constraints: removal of F3 prevented recording directly over the stimulation target, while the low-density montage limited spatial resolution, necessitated broad ROI definitions, and constrained anatomical interpretation of sensor-level PLV effects [64–67]. Higher-density EEG, source-space analyses, and connectivity measures less sensitive to zero-lag coupling could help clarify tSMS-related cortical interactions.

The relatively small sample size and marked sex imbalance also limit the generalizability of our findings, highlighting the need for larger, more balanced samples to investigate inter-individual variability, predictors of responsiveness, and potential sex-related differences. Repeated-session protocols could help determine whether cumulative stimulation produces more sustained electrophysiological changes and whether these translate into measurable behavioral effects. Such investigations would also help establish the potential translational relevance of left DLPFC tSMS for clinical populations with alterations in prefrontal function and its interactions with other cortical regions. However, given that the present study involved healthy participants and did not demonstrate behavioral improvement, clinical implications remain preliminary and require direct evaluation using neurophysiological measures and clinically relevant behavioral outcomes.

## 5. Conclusion

A single session of left DLPFC tSMS can modulate frontal spectral activity both during stimulation and during a subsequent cognitive task, as well as phase synchronization between frontal and centroparietal recording regions during stimulation, without significantly affecting divergent-thinking performance. These findings highlight the potential of prefrontal tSMS to modulate distributed electrophysiological dynamics, although their underlying mechanisms and functional significance remain to be established.

## Funding

This research did not receive any specific grant from funding agencies in the public, commercial, or not-for-profit sectors.

## Acknowledgements

The authors thank Fátima Villamayor-Vega and Sara Fernández-Canosa for their assistance as independent raters of AUT originality scores. The authors also thank Lara Ruiz Álvarez for her advice on AUT scoring procedures, and Manuel Nieto Díaz for helpful comments on the statistical approach.

## CRediT authorship contribution statement

Eva Díez-Rodríguez: Conceptualization, Methodology, Software, Formal analysis, Investigation, Data curation, Visualization, Project administration, Writing – original draft, Writing – review and editing.

Daniel Jimeno-Huete: Software.

Antonio Oliviero: Conceptualization, Methodology, Supervision, Project administration, Writing – review and editing.

Vanesa Soto-León: Conceptualization, Methodology, Validation, Supervision, Project administration, Writing – review and editing.

## Declaration of competing interests

AO declares that he is cofounder of Neurek SL, a spin-off of the Foundation of the Hospital Nacional de Parapléjicos. He is also listed as an inventor on patent P201030610. The remaining authors declare no competing interests.

## Declaration of generative AI and AI-assisted technologies in the manuscript preparation process

During the preparation of this work, the authors used ChatGPT to support language editing and the condensation of manuscript text to meet word-limit requirements. After using this tool, the authors reviewed and edited the content as needed and take full responsibility for the content of the published article.

## Notes

### Competing Interest Statement

AO declares that he is cofounder of Neurek SL, a spin-off of the Foundation of the Hospital Nacional de Paraplejicos. He is also listed as an inventor on patent P201030610. The remaining authors declare no competing interests.

