## Supplementary Methods for "Transcranial static magnetic field stimulation over the left dorsolateral prefrontal cortex modulates EEG spectral activity and phase synchronization without detectable changes in divergent thinking"

#### S1. Alternative Uses Task administration and scoring

Object names were presented on screen, and participants typed their responses in a text box. Before the experimental trials, participants received practice examples and were informed that common uses and multiple responses describing the same general use would be excluded. Two sets of objects were used across the pre- and post-stimulation assessments. Set A comprised a shoe, a newspaper, and an umbrella. Set B comprised a chair, a brick, and a paperclip.

Fluency. For each object, fluency was scored as the number of valid, distinct uses generated. Repeated or redundant responses, common uses, incomprehensible responses, and responses that did not describe a plausible use of the object were excluded.

Originality. Responses were compiled into a common list, and duplicates were removed before rating. Four judges, blinded to stimulation condition, independently rated each distinct response on a five-point scale. Judges considered both the rarity and the novelty of each use, with higher scores indicating greater originality. Ratings were averaged across judges. For each participant, mean originality was calculated for each object and then averaged across objects. Inter-rater reliability was assessed using the intraclass correlation coefficient for the mean of the four judges' ratings.

Flexibility. Flexibility was scored as the number of distinct semantic categories represented in a participant's responses. ChatGPT-4 was used to generate an initial set of candidate category labels. Trained human raters reviewed and refined these labels, defined the final categories, and made all scoring decisions.

Composite score. Fluency, flexibility, and originality scores were standardized across participants. Each participant's composite score was the mean of the three standardized scores.

#### S2. Additional baseline measures

The computerized sustained-attention task comprised 320 trials divided into four blocks, with a short break between the second and third blocks. On each trial, a black circle appeared in the center of a white screen after a randomly varying interstimulus interval of 1,000–2,500 ms. Participants were instructed to press a response button as quickly as possible when the circle appeared. Reaction time and accuracy were recorded. Mean reaction time was the primary index of sustained attention.

The Spanish versions of the State–Trait Anxiety Inventory and the NEO Five-Factor Inventory were scored according to their respective manuals.

**S3. EEG preprocessing**

Continuous EEG data were filtered using finite impulse response filters. Bad channels were identified with clean_rawdata using a flatline duration criterion of 5 s, a channel correlation threshold of 0.80, and a line-noise criterion of 4. For artifact subspace reconstruction, the burst criterion was set to 20 and the window criterion to 0.25, with rejection of contaminated segments enabled. Following independent component analysis, components were rejected if ICLabel assigned a probability of at least 0.70 to any of the following classes: muscle, eye, heart, line noise, or channel noise. Original sample positions were retained when assigning EEG data to experimental intervals. Discontinuities introduced by artifact rejection were preserved so that spectral estimation, band-pass filtering, and phase extraction could be performed separately on contiguous segments.

**S4. Spectral analysis and parameterization**

For conventional band-power estimation, Tukey's 1.5 × interquartile-range criterion was applied to $\log_{10}$-transformed power spectral density (PSD) values separately for each electrode, frequency bin, and experimental interval, pooling observations from both stimulation groups. Spectra with at least 76 of the 80 frequency bins valid were eligible for interpolation. Missing values between valid bins were interpolated linearly in linear power units. Missing values at either end of the spectrum were replaced with the nearest valid value. Spectra with more than four missing bins were excluded from band-power estimation. For each complete spectrum, absolute band power was calculated by summing the linear PSD values within each band and multiplying by the frequency-bin width (0.5 Hz).

FOOOF (version 1.1.1) was fitted to the original PSD estimates over 2–40 Hz. In fixed aperiodic mode, the $\log_{10}$ power spectrum was modeled as the sum of an aperiodic background and a set of Gaussian peaks.

$$P(f) = L(f) + \sum_{n=1}^{N} G_{n(f)}$$

Here, $P(f)$ denotes the modeled $\log_{10}$ power spectrum at frequency $f$ in Hz. In fixed mode, the aperiodic component is $L(f) = b - \chi \log_{10}(f)$, where b is the offset and $\chi$ is the aperiodic exponent. The slope of the aperiodic component in log–log coordinates is therefore $-\chi$. Each $G_n(f)$ represents the nth fitted Gaussian peak, and $N$ is the total number of fitted peaks.

Peak-width limits were set to 1–12 Hz, the maximum number of peaks was six, and the peak threshold was 2. For each ROI, the offset and exponent were averaged across electrode-level fits meeting the inclusion criterion of $R^2 \geq 0.70$. Example fits and distributions of fit-quality metrics are shown in Supplementary Fig. S1.

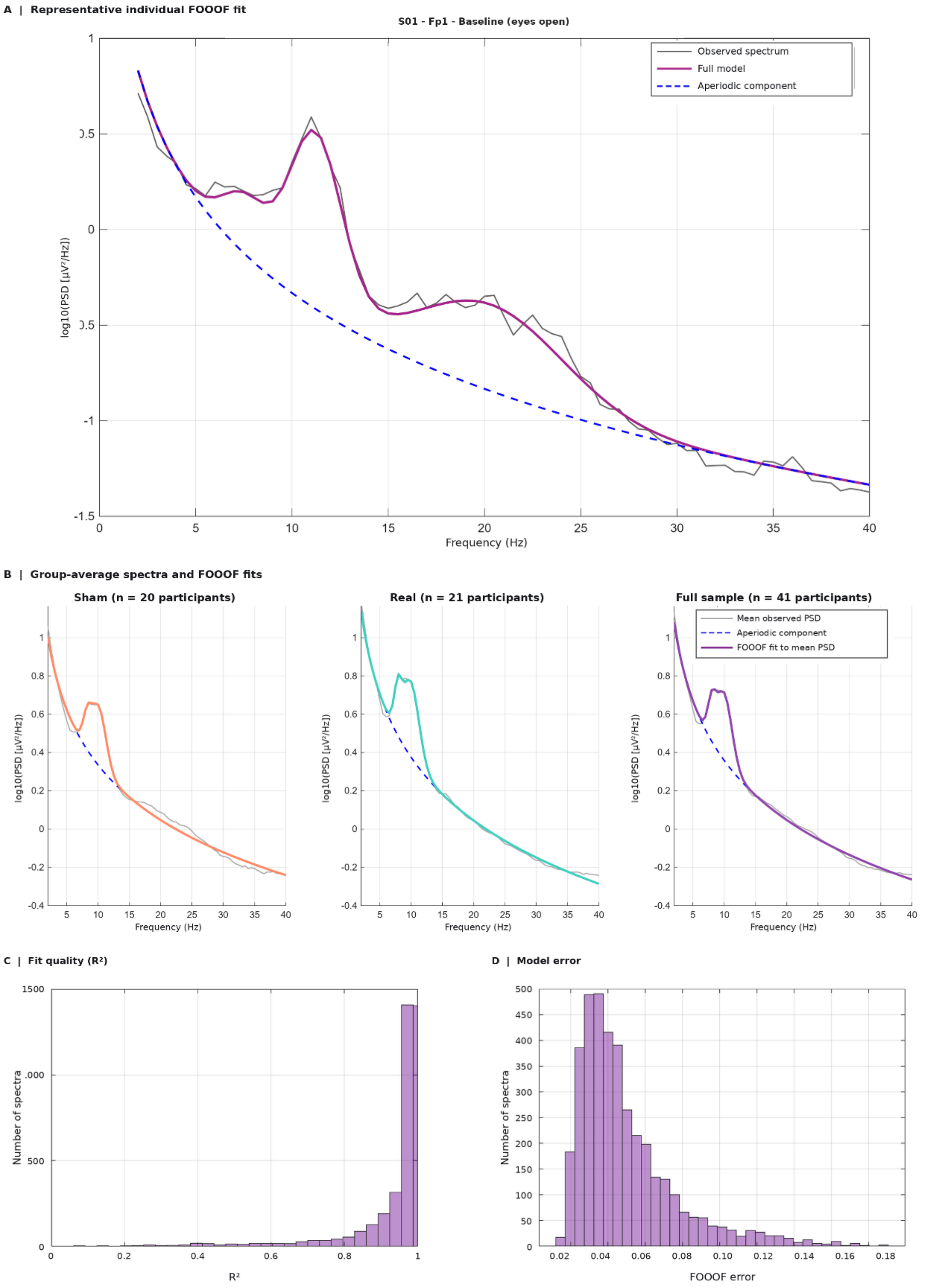

**Supplementary Figure S1. FOOOF spectral fits and fit quality across EEG spectra.** (A) Example fit for participant S01 at electrode Fp1 during the eyes-open baseline. The observed power spectrum (gray), full FOOOF model (magenta), and fitted aperiodic component (dashed blue) are shown over the 2–40 Hz fitting range. (B) Mean power spectra for the sham group ($n$ = 20), real tSMS group ($n$ = 21), and full sample ($n$ = 41). A separate FOOOF model was fitted directly to each mean spectrum. Solid colored lines show the full fits, and dashed blue lines show their aperiodic components. (C) Distribution of $R^2$ values across fitted spectra. Fits with $R^2 \geq 0.70$ were retained for the aperiodic analyses. (D) Distribution of FOOOF model error across spectra. In both histograms, the vertical axis indicates the number of spectra.

### S5. Phase synchronization

Each continuous EEG segment was filtered separately within each frequency band using a fourth-order Butterworth band-pass filter applied in the forward and reverse directions. Instantaneous phase was extracted from the analytic signal using the Hilbert transform. To limit edge effects, three cycles at the lower cutoff frequency of the band were discarded from each end of the segment. A segment was retained if at least five usable cycles remained after trimming. A minimum of 15 s of effective EEG data was required per frequency band and experimental interval.

For electrodes $x$ and $y$, the phase-locking value (PLV) was calculated separately for each frequency band and experimental interval from the instantaneous phase differences across valid samples.

$$PLV = \left| \left(\frac{1}{n}\right) \sum_{t=1}^{n} e^{i\left(\phi_{x(t)} - \phi_{y(t)}\right)} \right|$$

Here, n is the total number of valid samples for the electrode pair, frequency band, and experimental interval under analysis. The index t runs from 1 to $n$ over these samples. The terms $\varphi_x(t)$ and $\varphi_Y(t)$ denote the instantaneous phases at electrodes $x$ and $y$, respectively. PLV ranges from 0 to 1, with higher values indicating a more consistent phase difference over time.

PLV was first calculated for each electrode pair. For intra-regional synchronization, the PLVs of all unique pairs of distinct electrodes within an ROI were averaged to obtain one value for that ROI. For inter-regional synchronization, the PLVs of all electrode pairs with one electrode in each of two distinct ROIs were averaged to obtain one value for that ROI pair. Fisher's z transformation was applied after averaging the electrode-pair PLVs.
